# Genetic parameters and genotype-by-diet interactions for growth traits in Australian black soldier fly larvae: Implications for selective breeding

**DOI:** 10.64898/2026.03.14.711759

**Authors:** K. B. Gowda, S. Septriani, D. B. Jones, D. R. Jerry, C. Tedder, K. R. Zenger

## Abstract

Black soldier fly larvae (*Hermetia illucens*, Diptera: Stratiomyidae; BSFL) efficiently bio- convert organic waste into high-value protein, which has significant potential in domesticated animal feed formulations. BSFL growth and bioconversion potential can be enhanced through selective breeding, which requires accurate estimates of genetic parameters and knowledge of genotype-by-diet (G × D) interactions. However, comprehensive knowledge of G × D interactions is limited, and reports of genetic parameters are sparse across genetic strains and production environments globally. This study estimated heritabilities, dominance effects and genetic correlations for BSFL growth traits and quantified G × D interactions. Phenotypes of 2,097 fifth-instar larvae reared on three diets were recorded, including larval body weight (LBW), length (LL), width (LW), and surface area (LSA). All larvae were genotyped using a custom 6K Allegro SNP panel. Genetic parameters and G × D interactions were estimated by fitting additive and additive-dominance models. Heritabilities for growth traits were low across diets (0.04-0.14), with diet-specific estimates ranging from low to moderate (0.08- 0.37). Dominance effects were significant across the traits (0.10-0.19), and genetic correlations were high among growth traits (>0.79), except between LW and LL (0.47). G × D interactions were moderate across diets (−0.06-0.46). Results suggest that moderate to high genetic gain is achievable over a long-term breeding program, given the genetic basis of growth traits and BSF’s short generation interval (38-45 days). However, G × D interactions must be considered, either through combined or diet-specific selection strategies, and the significant dominance effects suggest heterosis could accelerate improvement.

## 1. Introduction

The Black soldier fly (BSF) (*Hermetia illucens*, Diptera: Stratiomyidae) is a promising insect species for sustainable farming within the emerging sector of insects as human food and animal feed (van Huis, 2013; Wang and Shelomi, 2017). With a reported bioconversion rate of 0.2-55.8%, BSF can convert a wide range of organic waste into protein and organic fertiliser, thereby contributing to the circular bioeconomy (Siddiqui *et al*., 2024). BSF larvae (BSFL) are distinguished by their high crude protein content (36-65%), variable fat content (4.6-38.6%), and the presence of antimicrobial peptides (Hua *et al*., 2019; Mohan *et al*., 2022; Vogel *et al*., 2018). These attributes facilitate their reintroduction into food production systems, particularly as ingredients for aquaculture and livestock feed. Early projections suggest that the industry holds strong growth potential, with an estimated market value of US$4 billion by 2033 and a compound annual growth rate (CAGR) of 33.7% (Bukchin-Peles *et al*., 2025). To realise this, various efforts are underway to industrialise BSFL production, including automation, improvements in production infrastructure and environmental control, and the optimisation of breeding, nutrition, and grow-out management.

Specifically, the commercial production of BSFL can be further enhanced through the genetic improvement of traits such as growth, reproduction, and feed conversion efficiency via selective breeding (Eriksson and Picard, 2021; Gowda *et al*., 2025). Selective breeding is a well-established approach widely applied in livestock, plants, aquaculture and even in insects (e.g., the honeybee, *Apis mellifera*) (Crossa *et al*., 2017; Hayes *et al*., 2009; Hoppe *et al*., 2020; Jerry *et al*., 2022). It aims to enhance productivity by selecting individuals with high genetic merit to contribute to the next generation. To date, only a few studies have applied mass selection in BSFL, a selection approach in which parents are chosen based on superior phenotypic performance, achieving improvements such as a 39% increase in larval weight after 16 generations (Facchini *et al*., 2022), a 24-30% increase in fecundity after 32 generations (Shrestha *et al*., 2025) and enhanced thermal tolerance after 9 generations (Feng *et al*., 2025; Ma *et al*., 2024), demonstrating measurable gains. Global BSF populations are genetically diverse, and understanding their performance across production environments is essential for the true industrialisation of the species. However, current breeding efforts are constrained by the lack of accurate estimates of genetic parameters across global BSFL stocks and production environments, which are essential for designing effective breeding programs.

Genetic parameters such as heritability, genetic correlations, and breeding values are fundamental for understanding variation in production traits and predicting responses to selection (Falconer, 1960). Heritability quantifies the proportion of phenotypic variation due to additive genetic effects; genetic correlations explain the genetic relationship among traits and potential correlated responses, and breeding values estimate the genetic merit of individuals for a specific (set of) trait to guide selection decisions. Accurate estimation of these parameters is crucial for designing efficient breeding programs for BSFL. To date, only a few studies have estimated genetic parameters for BSFL populations from Europe, Africa and North America, with heritability estimates ranging from 0.17 to 0.78 for larval and pupal size and around 0.37 for developmental time (Bouwman *et al*., 2022; Generalovic *et al*., 2025a; Hull *et al*., 2024; Zaalberg *et al*., 2025). Most of these studies relied on pedigree-based full-sib or half-sib designs, in which families were reared separately; such designs tend to inflate estimates because non-additive genetic and common environmental (rearing tray) effects cannot be separated effectively. While Hull *et al*. (2024) used SNP markers to estimate genetic parameters for the South African population, their analysis was based on a relatively small sample size (n=146), with SNP-based heritability for individual larval mass estimated at 0.179-0.184. Although these estimates had relatively small standard errors (0.029-0.033), larger-scale studies using genome-wide SNP data, combined with rearing individuals under standardised and well-characterised environmental conditions to minimise confounding effects, are likely to yield more accurate and unbiased parameter estimates. However, no such estimates are currently available, particularly for the Australasian lineage, which is genetically distinct from other global stocks and has reduced genetic diversity due to its introduction history and subsequent isolation (Generalovic *et al*., 2025b; Kaya *et al*., 2021). Genetic differentiation has also been observed among Australian populations, with pairwise *F*_ST_ values ranging from 0.050 to 0.276 (Davidson *et al*., 2026; in revision).

Environmental variation can influence genotype performance, resulting in genotype-by-environment interactions (G **×** E), whereby genotypes respond differently when they are exposed to different environments (Falconer, 1960). These interactions influence phenotypic plasticity, selection objectives and the expected response to selection within breeding programs (Saltz *et al*., 2018) and are well documented in animal, plant, and aquaculture production systems (Des Marais *et al*., 2013; Jerry *et al*., 2022; Silva Neto *et al*., 2024). In quantitative genetics, G × E interactions are commonly assessed using genetic correlations for the same trait measured across environments, with lower genetic correlations indicating greater genotype re-ranking and may necessitate environment-specific selection. Consequently, assuming the absence of G × E interactions may reduce selection efficiency when breeding and production environments differ. Despite their importance in managed production systems, G × E interactions have received little attention in managed insects, with limited evidence in honey bees (Gowda *et al*., 2025; Meixner *et al*., 2015).

BSFL can be raised on various organic substrates, which differ in their nutrient composition, with production often depending on locally available organic waste (Tognocchi *et al*., 2024). Understanding how BSF genotypes or strains interact with different diets in a commercial production setup is essential for optimising the breeding programs (Sandrock *et al*., 2022). In BSF production systems, growth rate and final proximate composition primarily depend on nutritional factors in their diet and hold significant economic value. Therefore, resolving interactions between genotype and diet (G × D) within the BSF production paradigm is essential for identifying strains that perform optimally on specific diets or substrates, or for breeding BSF strains that perform consistently across different diets. To date, two studies have investigated the G × D interaction in BSF and explained trade-offs between genetic strains and production traits (Generalovic *et al*., 2025a; Sandrock *et al*., 2022). However, these studies were limited to comparing distinct pre-selected genetic lines rather than a diverse, mixed breeding population.

The present study addresses key knowledge gaps by estimating heritabilities and genetic correlations of key growth traits in Australian BSFL using genome-wide SNP data. It also examines G × D interactions, thereby providing baseline information to support the development of selective breeding programs and to strengthen the farming of BSFL and its contribution to the circular bioeconomy.

## 2. Materials and Methods

### 2.1 Ethics statement

Research involving BSFL in Australia does not require ethical approval.

### 2.2 Base population

The base population for this study was established using BSFL collected from 10 geographically distinct locations across Australia between November 2023 and February 2024, with the aim of capturing broad genetic diversity and local environmental adaptations. Pairwise *F*_ST_ estimates among these source populations ranged from 0.050 to 0.276, while population genetic analyses of these source populations, including principal component analysis (PCA), are described in detail elsewhere (Davidson *et al*., 2026; in revision; supplementary figure 1). At each location, at least 1,000 wild late-instar (instar 5) larvae were collected and transported to the BSF rearing facility at James Cook University (JCU), Townsville, Australia.

Upon arrival, BSFL from each population were reared separately in insect cages (BugDorm- 6E610; 60 × 60 × 60 cm, 0.216 m³, MegaView Science Co., Ltd., Taichung, Taiwan) to maintain their genetic integrity and establish sizeable breeding populations. The larvae were allowed to pupate, emerge as adults, and mate. Each cage was equipped with a water source for adult hydration and egg collectors to harvest eggs. Egg clutches were collected, hatched, and reared on a commercial chicken feed diet containing 15.5 % protein, with moisture maintained at 60-70%.

To establish a genetically diverse experimental population, these initial source populations were subsequently transferred to the commercial breeding facility in Brisbane. Pupae from all populations were pooled and randomly allocated to six 1 m³ cages, each stocked with 20,000 pupae to initiate outbreeding. This approach was adopted to simulate commercial breeding colonies, which are expected to comprise genetically diverse breeding populations rather than geographically isolated or predefined strains. Husbandry across breeding, nursery, and grow-out stages followed the facility’s standard operating procedures, with the company’s commercial diet used throughout all production phases. The outbred stocks underwent three further generations under these controlled conditions. In each generation, pupae from all cages were pooled and randomly redistributed to ensure consistent genetic mixing before the experiment began.

Throughout both JCU and commercial phases, rearing was conducted under standardised environmental conditions, including a 12:12 h light-dark cycle with UV lighting (BSF-4C- 200-3030 LED light, SPR Ag Tech) to support mating, temperature maintained at 27-30 °C, and relative humidity controlled between 60-70%.

### 2.3 Experimental design and setup

#### 2.3.1 Breeding design

The experiment was conducted at a commercial R&D facility, using the third-generation outbred base population. Adults from six cages (stocked with 20,000 pupae each) were allowed to mate and oviposit under a mass breeding design using egg collectors. Eggs were collected daily, and the 53 egg clutches (∼1.188 g total, representing putative full-sib families) collected on the morning of the fourth day of the collection cycle were used for the experiment. The eggs were incubated in a single tray containing a nursery diet of chick starter, soy okara, mill run, and water (5:1:1:7). The climate-controlled chamber was maintained at 29 °C and 60% relative humidity. Eggs hatched within three days, and neonates were kept in the same nursery tray until day 7 post-egg collection.

#### 2.3.2 Diets and larval allocation to grow out

Three experimental diets were used in this study: soy okara (SYK) and brewer’s spent grain (BSG), which were sourced from food processing industries and frozen upon arrival. Fruit and vegetable waste (FVW) was collected from local shops and markets, ground into small particles, pooled and similarly stored at -18 °C as a single batch. Proximate analysis of all three diets was conducted, with crude protein determined by the Kjeldahl method, crude fat quantified using Soxhlet extraction, and ash by incineration (Greenfield *et al*., 2003) (Supplementary Table 1). All diets were thawed 24 h before the start of the grow-out phase and added to trays at a rate of 4 kg per tray. Feed moisture content was adjusted and maintained between 60 and 70%.

On day 7, neonates were separated from the frass, counted, and randomly distributed into nine trays (Insect breeding box, 145 mm; Beekenkamp Verpakkingen B.V., Netherlands) with ∼1551 neonates/tray (average weight 23.45 mg). Trays were assigned to one of three diets in triplicate: SYK, BSG, or FVW. The trays were arranged in two vertical columns (randomly stacked) and covered with mesh to prevent larval movement between trays. All trays were housed in a 1 m^3^ cage mounted on a rack at 1.2 m above ground level. The temperature was maintained at an average of 27 °C, and relative humidity was kept at 70% throughout the grow-out phase.

### 2.4 Phenotyping

Larvae from all the diet treatments and trays were sampled at the fifth instar stage on days 13, 14 and 15 post-egg collection (supplementary Table 2). The recorded phenotypic traits were larval length (LL), width (LW), surface area (LSA), and colour, obtained from digital images. Prior to imaging, larvae were immobilised by placing them at -18 °C for 10 min. Images were captured using a Sony a6500 camera (Sony Australia Ltd.), positioned 30 cm above the subject. Each larva was placed individually into wells carved into a white Teflon board, housed within a light box, and accompanied by a scale ruler and colour palette (see supplementary figure 2). After imaging, individual larval body weight (LBW) was also measured using a precision digital balance (IC-PX124, Ohaus Australia). Each larva was then bisected and preserved in chilled ethanol for subsequent genotyping. Digital images were analysed manually for LL, LW, LSA, and colour (RGB scale) using ImageJ software (Schneider *et al*., 2012), based on key morphological landmarks. Larval sex could not be determined morphologically at this developmental stage and was therefore not considered in the statistical models (section 2.7).

### 2.5 Genotyping and quality control of SNPs

Genomic DNA was extracted from the head tissue of preserved BSFL samples using the automated high-throughput Chemagic™ 360 magnetic bead extraction system (PerkinElmer Inc.), following the manufacturer’s protocol for the CMG-723 tissue kit. Briefly, 20 mg of tissue was incubated with 6 μL of Proteinase K and 200 μL of lysis buffer at 56 °C for 1 h. After incubation, 150 μL of magnetic beads and binding buffer were added to each sample. Magnetic separation was used to capture the DNA-bound beads, followed by four wash steps. DNA was eluted in 150 μL of elution buffer. DNA quality was assessed by 0.8 % agarose gel electrophoresis in 1x TAE buffer, and concentration was measured using the QuantiFluor™ dsDNA Dye System (Promega). Samples meeting the quality criteria (minimum 20 ng/μL in 20 μL volume) were submitted to the Australian Genome Research Facility (AGRF Ltd.) for SNP genotyping using the Allegro® Targeted Genotyping V2 (Tecan Genomics, Switzerland) platform with a custom 6K SNP panel, which was designed and validated in-house for Australian BSF (Davidson *et al*., 2026; in revision). SNPs were called using the Genome Analysis Toolkit (*GATK v4.2.0.0*) best-practices pipeline for targeted sequencing, using the *Hermetia illucens* (iHerIll2.2.curated.20191125, GenBank accession No. GCF_905115235.1; (Generalovic *et al*., 2021)) reference genome for read alignment and variant identification. SNPs were filtered using *PLINK v1.9* (Purcell *et al*., 2007) based on a minor allele frequency (MAF) threshold of 0.5%, an individual call rate of 90% and a minimum SNP call rate of 90%.

### 2.6 Population structure and sibship analysis

Population structure was evaluated using PCA implemented in *PLINK v1.9* (Purcell *et al*., 2007). PCA was conducted on the filtered SNP dataset (see Section 2.5), and population structure was visualised by plotting the first two principal components in *ggplot2* package in R (Wickham and Sievert, 2009), with individuals grouped by diet to assess the distribution of genetic groups across the three diet treatments (SYK, BSG and FVW).

Sibship analysis was performed using *COLONY* (Jones and Wang, 2010) to infer family clusters among individuals. The analysis used 2,083 individuals genotyped at 1,200 informative SNPs (from the filtered genotypic data, Section 2.5), assuming a dioecious diploid species with inbreeding. A full-likelihood method was used with a long run and high precision, with no clone or sibship specified. The inferred family clusters were subsequently assessed for their distribution across three diet treatments.

### 2.7 Statistical analysis

#### 2.7.1 Phenotypic data curation

The normality of phenotypic data was assessed using the *PROC UNIVARIATE* procedure in SAS 9.4. No major deviations from normality were observed, and the data were considered approximately normally distributed. Prior to this assessment, descriptive statistics for the complete dataset were generated using the *PROC MEANS* in SAS 9.4.

#### 2.7.2 Non-genetic factors

Diet (SYK, BSG and FVW), sampling day (13, 14 or 15), trays nested within diets, and larval colour (a potential indicator of physiological/developmental changes) were evaluated as non-genetic factors. Their effects on growth traits were analysed using a generalised linear model (GLM), in which these variables were specified as fixed effects. The analysis was carried out using the *proc glm* procedure in SAS 9.4 to assess the statistical significance of each factor. Factor-specific effects were estimated using least-squares means from the final model, and comparisons among levels were performed using the Tukey-Kramer adjustment. Type III mean squares were used to assess each factor’s contribution, and the coefficient of determination (R^2^) was used to quantify the proportion of phenotypic variation explained by the model. The initial model incorporated all main effects and interaction terms, with only statistically significant interactions retained in the final model. Additionally, to account for population structure, PCs 1-4 and genetic clusters derived from PCA were evaluated as fixed effects. However, neither the PCs nor genetic clusters had significant effects on growth traits (*p* > 0.05) and were therefore not considered in the final model.

#### 2.7.3 Estimation of genetic parameters

Genetic variance and covariance components for growth traits (LBW, LL, LW and LSA) were estimated in ASReml-R version 4.1 (Butler *et al*., 2017) via the residual maximum likelihood (REML) approach. Two alternative models were evaluated.

Model 1: Additive model

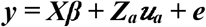

Where ***y*** represents the vector of observed phenotypes, and β denotes the vector of fixed effects, including the overall mean, colour (covariate), diet, sampling day, and tray. The term ***u_a_*** corresponds to the vector of random additive genetic effects, and ***e*** represents the vector of random residual effects. The matrices ***X*** and ***Z_a_*** are the incidence matrices linking observations to fixed and additive genetic effects, respectively.

The random effects were assumed to follow multivariate normal distributions, with 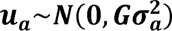, where 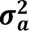 is the additive genomic variance, and ***G*** is the genomic relationship matrix (GRM) constructed from genome-wide SNPs using the snpReady (Granato *et al*., 2018) package in R, following the method of VanRaden (2008). Residual effects were assumed to follow 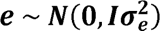, where 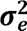 is the residual variance, and ***I*** is the identity matrix.

The heritability (***h^2^***) for each trait was calculated as

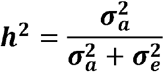

Model 2: Additive-dominance model

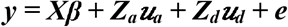

This model extends Model 1 by adding an additional dominance component. Here,**u*_d_*** represents the random dominance genetic effects, and ***Z_d_*** maps observations to these effects. The dominance relationship matrix was constructed using the method described by Vitezica *et al*. (2013). All other parameters are as described for the additive model.

The genetic and phenotypic correlations of growth traits (LBW, LL, LW and LSA) were estimated by fitting the bivariate additive-dominance models in ASReml-R version 4.1 (Butler *et al*., 2017).

Genetic correlations were estimated as:

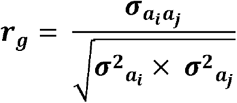

Where 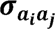 denotes the additive genomic covariance shared by the two traits and 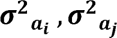 represents the additive genomic variances of the respective traits.

Phenotypic correlations were estimated as:

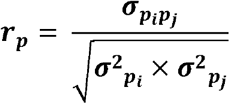

Where, 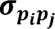 denotes the phenotypic covariance shared by the two traits and 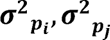 represents the phenotypic variances of the respective traits.

#### 2.7.4 Quantification of G x D

G × D was quantified by a multi-trait additive model, as the additive dominance model failed to converge, in which the same trait measured in three different diets was treated as three independent traits. This was implemented as a multivariate animal model in ASReml-R version 4.1 (Butler *et al*., 2017):

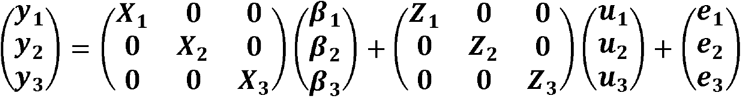

Where **y_1_**, **y_2_** and **y_3_** are the same trait measured in diets SYK, BSG, and FVW, respectively. Fixed and random effects were specified as in the additive model (section 2.7.3). Genetic correlations in trait expression across diets were used to assess the magnitude of G × D.

Reaction norms were generated by plotting genomic estimated breeding values (GEBVs) from the multi-trait additive model above for the top 5% of individuals across the three diets, with each individual’s genetic values connected across diets to visualise genotype re-ranking.

## 3. Results

### 3.1 Dataset and descriptive statistics

The dataset comprised 2,097 individual larvae collected across three diets (SYK: 973, BSG: 560, and FVW: 564), with growth traits measured and genotypes generated using the 6K Allegro SNP panel. Differences in sample numbers among diets arose from sampling logistics and subsequent genotyping and quality-control processes, not from differences in the initial experimental allocation. Of these, 2,083 samples passed the primary quality-control filters, with a sample call rate of 90%, SNP call rate of 90%, and a minor allele frequency threshold of 0.5%, resulting in a final dataset of 4,680 SNPs for 2,083 individuals.

The descriptive statistics (mean ± SD) for BSFL growth traits are presented in Table 1. The coefficient of variation ranged from 18–30% for LBW across diets, whereas LSA ranged from 17–27% across diets (Supplementary Table 3). The mean LBW was highest for larvae reared in SYK, followed by BSG and FVW (Table 1). Overall, growth traits varied substantially across diets, with SYK producing the largest larvae and FVW the smallest. The traits were not strictly normally distributed (Figure 1); however, they were assumed to be approximately normal. After fitting the model in ASReml-R, residual plots and Q-Q plots were inspected to verify this assumption, and no major deviations were observed.

**Figure 1:**
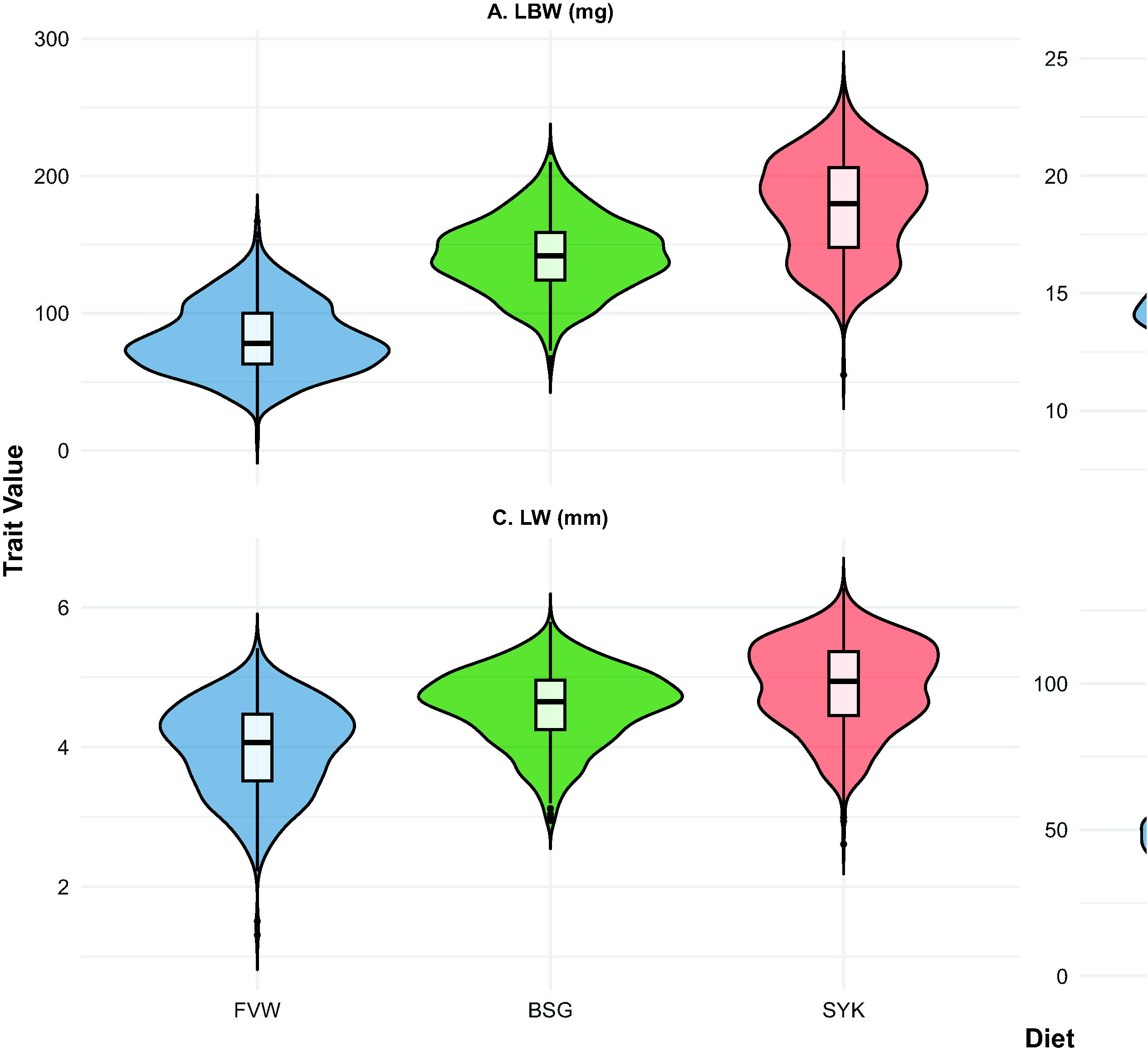
Distributions of growth traits, including larval body weight (LBW), length (LL), width (LW) and surface area (LSA), across diets (SYK: soy-okara, BSG: brewers’ spent grain, FVW: fruit and vegetable waste).

**Table 1:** Mean (± standard deviation) of larval body weight (LBW), length (LL), width (LW) and surface area (LSA) across diets (SYK: soy-okara, BSG: brewers’ spent grain, FVW: fruit and vegetable waste) and overall; N denotes the number of records.

| Diet | N | LBW, mg | LL, mm | LW, mm | LSA, mm<sup>2</sup> |
| --- | --- | --- | --- | --- | --- |
| SYK | 973 | 177.11±35.97 | 19.02±2.04 | 4.87±0.62 | 79.60±18.49 |
| BSG | 560 | 141.85±26.60 | 17.80±1.75 | 4.57±0.53 | 69.49±12.48 |
| FVW | 564 | 81.82±25.33 | 14.17±1.74 | 3.97±0.65 | 47.58±12.86 |
| Overall | 2097 | 142.07±50.07 | 17.39±2.76 | 4.55±0.71 | 68.29±20.49 |

PCA analysis of the experimental population revealed distinct genetic clusters, with PC1 and PC2 explaining 15.39% and 13.31% of the total genetic variation, respectively (Figure 2). Individuals from the diet treatments were distributed across these genetic clusters, with substantial overlap among diets. Sibship analysis identified 64 unique family clusters across the 2,083 genotyped individuals, with family cluster sizes ranging from 1 to 148 larvae (Supplementary Table 4). Members from 41 of these family clusters were distributed across all three diet treatments.

**Figure 2:**
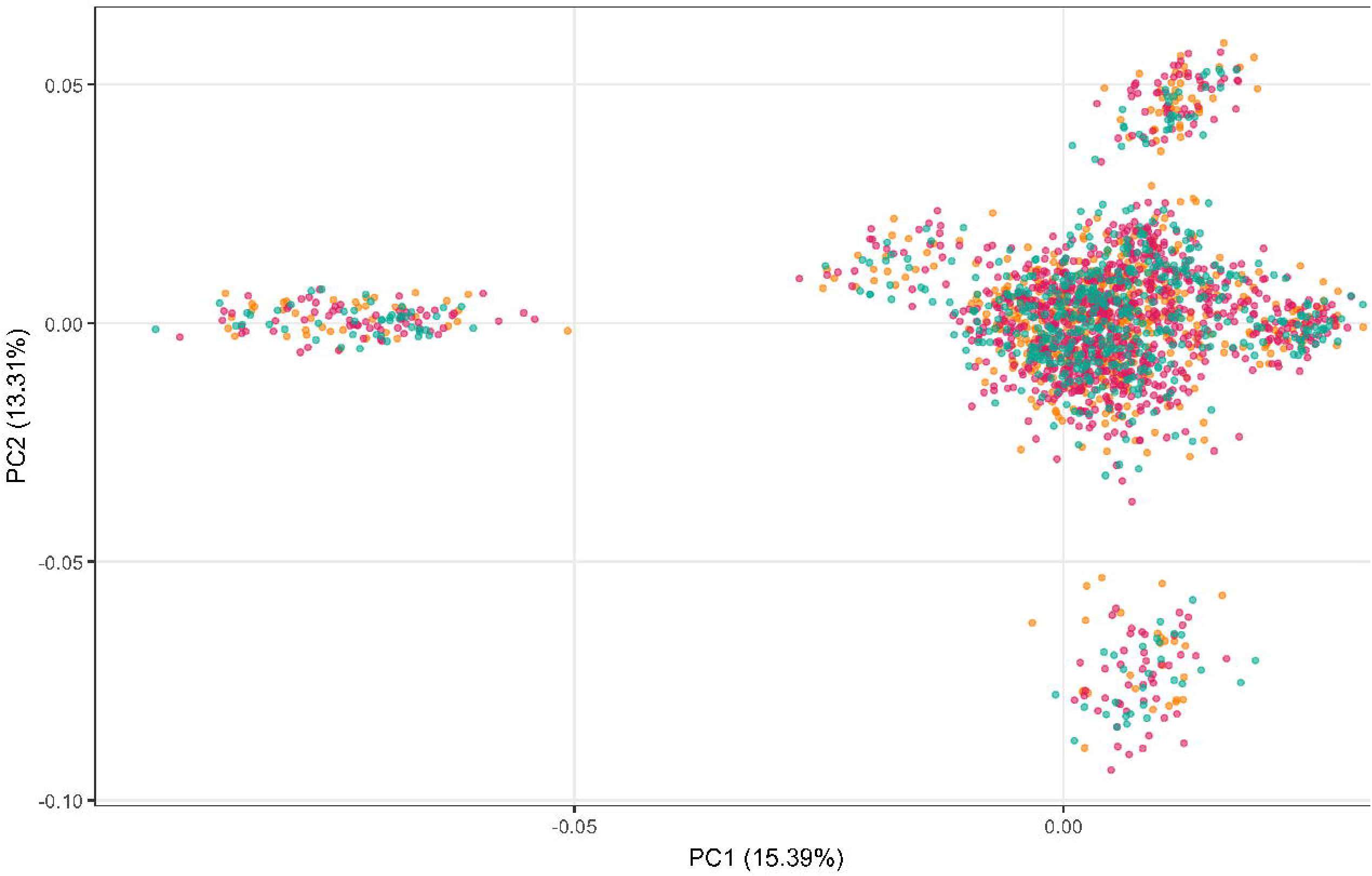
Principal component analysis (PCA) showing the distribution of the genetic groups across diet treatments (SYK: soy-okara, BSG: brewers’ spent grain, FVW: fruit and vegetable waste).

### 3.2 Effect of Non-genetic factors

The non-genetic factors considered in this study: diet, sampling day, trays within diets, and larval colour (as a covariate), had statistically significant effects on all BSFL growth traits (*p* < 0.05). We therefore retained all main effects in the final models. Collectively, these factors accounted for 65.75%, 57.82%, 34.65%, and 53.81% of phenotypic variation in LBW, LL, LW and LSA, respectively. Diet accounted for the largest proportion of variation across all growth traits. The least squares means (LS means) estimated from the model showed significant differences among diet treatments (Table 2), with Tukey-Kramer-adjusted pairwise comparisons indicating that all three diets differed significantly from each other (*p* < 0.05). Sampling day had relatively smaller effects, with day 14 generally differing from days 13 and 15 for LBW, LL and LSA, whereas no significant differences were observed among sampling days for LW. Differences among trays within diets were generally small, although significant differences were observed for some traits (Table 2). Type III mean squares from the model further showed that the diet effect was greater than other non-genetic effects across the growth traits (Table 3). Proximate analysis of the diets also revealed notable differences in nutrient composition (Supplementary Table 1). Lipid content ranged from 1.23% in FVW to 9.37% in BSG, protein content ranged from 7.3% in FVW to 21.3% in SYK, and ash content was relatively similar across diets (3.75-4.28%).

**Table 2:**
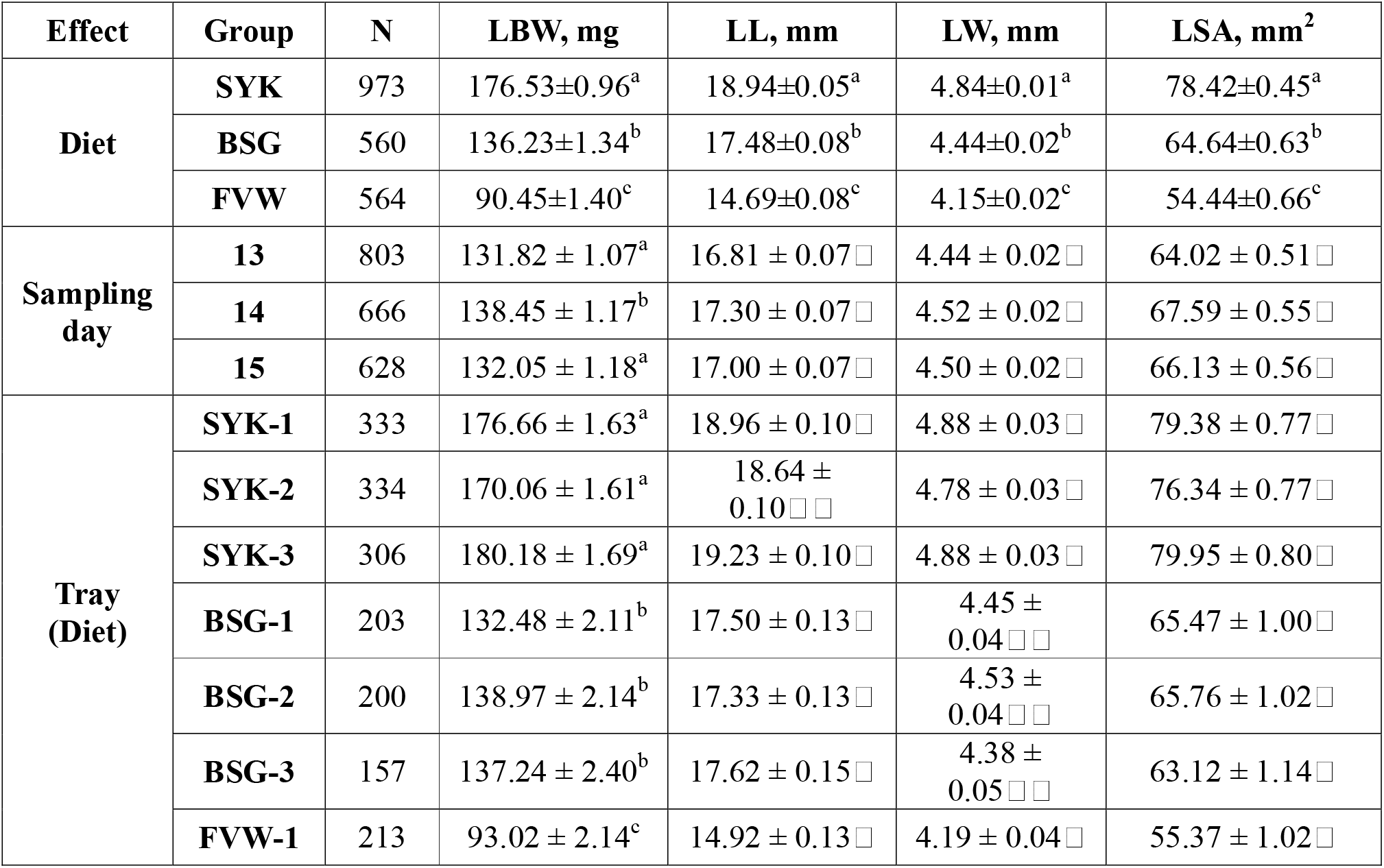
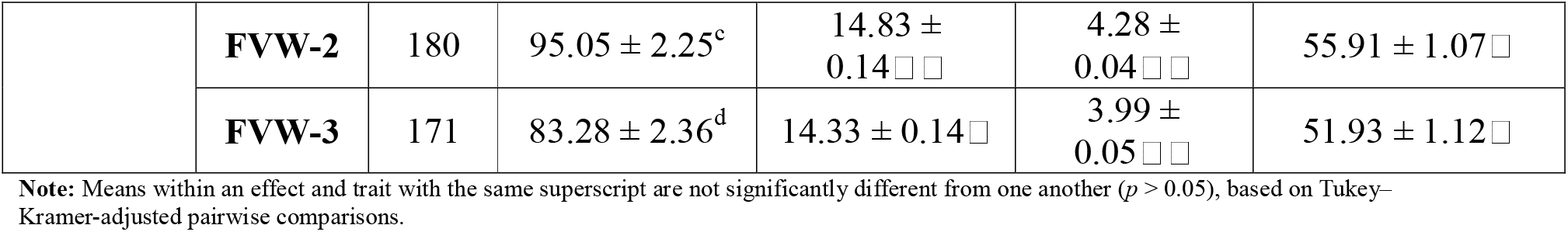
Number of observations (N) and least squares means (± standard error) for larval body weight (LBW), length (LL), width (LW) and surface area (LSA) by diet (SYK: soy- okara, BSG: brewers’ spent grain, FVW: fruit and vegetable waste), sampling day, and trays nested within diet.

| Effect | Group | N | LBW, mg | LL, mm | LW, mm | LSA, mm<sup>2</sup> |
| --- | --- | --- | --- | --- | --- | --- |
| Diet | SYK | 973 | 176.53±0.96<sup>a</sup> | 18.94±0.05<sup>a</sup> | 4.84±0.01<sup>a</sup> | 78.42±0.45<sup>a</sup> |
| | BSG | 560 | 136.23±1.34<sup>b</sup> | 17.48±0.08<sup>b</sup> | 4.44±0.02<sup>b</sup> | 64.64±0.63<sup>b</sup> |
| | FVW | 564 | 90.45±1.40<sup>c</sup> | 14.69±0.08<sup>c</sup> | 4.15±0.02<sup>c</sup> | 54.44±0.66<sup>c</sup> |
| Sampling day | 13 | 803 | 131.82 ± 1.07<sup>a</sup> | 16.81 ± 0.07 □ | 4.44 ± 0.02 □ | 64.02 ± 0.51 □ |
| | 14 | 666 | 138.45 ± 1.17<sup>b</sup> | 17.30 ± 0.07 □ | 4.52 ± 0.02 □ | 67.59 ± 0.55 □ |
| | 15 | 628 | 132.05 ± 1.18<sup>a</sup> | 17.00 ± 0.07 □ | 4.50 ± 0.02 □ | 66.13 ± 0.56 □ |
| Tray (Diet) | SYK-1 | 333 | 176.66 ± 1.63<sup>a</sup> | 18.96 ± 0.10 □ | 4.88 ± 0.03 □ | 79.38 ± 0.77 □ |
| | SYK-2 | 334 | 170.06 ± 1.61<sup>a</sup> | 18.64 ± 0.10 □ □ | 4.78 ± 0.03 □ | 76.34 ± 0.77 □ |
| | SYK-3 | 306 | 180.18 ± 1.69<sup>a</sup> | 19.23 ± 0.10 □ | 4.88 ± 0.03 □ | 79.95 ± 0.80 □ |
| | BSG-1 | 203 | 132.48 ± 2.11<sup>b</sup> | 17.50 ± 0.13 □ | 4.45 ± 0.04 □ □ | 65.47 ± 1.00 □ |
| | BSG-2 | 200 | 138.97 ± 2.14<sup>b</sup> | 17.33 ± 0.13 □ | 4.53 ± 0.04 □ □ | 65.76 ± 1.02 □ |
| | BSG-3 | 157 | 137.24 ± 2.40<sup>b</sup> | 17.62 ± 0.15 □ | 4.38 ± 0.05 □ □ | 63.12 ± 1.14 □ |
| | FVW-1 | 213 | 93.02 ± 2.14<sup>c</sup> | 14.92 ± 0.13 □ | 4.19 ± 0.04 □ | 55.37 ± 1.02 □ |
|  | **FVW-2** | 180 | 95.05 ± 2.25<sup>c</sup> | 14.83 ± 0.14□□ | 4.28 ± 0.04□□ | 55.91 ± 1.07□ |
|  | **FVW-3** | 171 | 83.28 ± 2.36<sup>d</sup> | 14.33 ± 0.14□ | 3.99 ± 0.05□□ | 51.93 ± 1.12□ |

**Note:** Means within an effect and trait with the same superscript are not significantly different from one another ( $p > 0.05$ ), based on Tukey–Kramer-adjusted pairwise comparisons.

**Table 3:** Mean squares and coefficient of determination (R^2^) from the fixed-effect model evaluating the source of variation for larval body weight (LBW), length (LL), width (LW) and surface area (LSA).

| Source of variation | DF | LBW | LL | LW | LSA |
| --- | --- | --- | --- | --- | --- |
| Colour | 1 | 159569.4* | 581.69* | 65.33* | 96653.7* |
| Diet | 2 | 1115733.36* | 2634.66* | 78.57* | 96491.79* |
| Tray (Diet) | 6 | 5901.11* | 16.69* | 1.92* | 798.02* |
| Sampling Day | 2 | 9509.7* | 43.43* | 1.37* | 2316.07* |
| Error | 2085 | 863.36 | 3.24 | 0.33 | 194.97 |
| $R^2$ (%) | | 65.75 | 57.82 | 34.65 | 53.81 |

\* ( $p$ -value <0.05); DF, degrees of freedom.

### 3.3 Genetic parameters

The heritability of the growth traits was estimated using models 1 and 2. Heritability estimates, corresponding variance components, and model fit statistics are summarised in Table 4 for the whole dataset and in Table 5 for the diet-specific analyses. Model 1, which included only the additive genetic effects, yielded higher estimates of additive genetic variance and heritability, ranging from 0.08 to 0.20 across all traits in the whole dataset. In contrast, Model 2, which incorporated both additive and dominance effects, yielded lower heritability estimates ranging from 0.04 to 0.14, indicating a decrease in additive genetic variance and residual variances when dominance effects were accounted for. Overall, heritability estimates ranged from low to moderate.

**Table 4:** Variance components, heritability (± standard error) and model fit statistics for larval body weight (LBW), length (LL), width (LW) and surface area (LSA) from the full dataset using additive (A) and additive + dominance (A + D) models.

| Trait | Model | Variance parameters |  |  |  | Ratios |  |  | Model fit statistics |  |  |
| --- | --- | --- | --- | --- | --- | --- | --- | --- | --- | --- | --- |
|  |  | A (Additive) | D (Dominance) | E (Residual) | P (Phenotypic) | h<sup>2</sup> (Additive) | d<sup>2</sup> (Dominance) | e<sup>2</sup> (Error) | LogLik | BIC | AIC |
| LBW | A | 182.44±32.00 | – | 711.63±25.12 | 894.08±33.97 | 0.20±0.03 | – | 0.79±0.03 | -8008.00 | 16031.29 | 16020.02 |
| | A + D | 117.88±29.21 | 167.89±27.66 | 575.70±23.58 | 861.48±33.22 | 0.14±0.03 | 0.19±0.02 | 0.66±0.03 | -7930.85 | 15884.61 | 15867.70 |
| LL | A | 0.29±0.07 | – | 2.98±0.10 | 3.27±0.10 | 0.09±0.02 | – | 0.91±0.02 | -2268.31 | 4541.89 | 4540.62 |
| | A + D | 0.13±0.07 | 0.33±0.08 | 2.76±0.10 | 3.22±0.10 | 0.05±0.02 | 0.10±0.02 | 0.85±0.02 | -2251.01 | 4524.94 | 4508.03 |
| LW | A | 0.03±0.00 | – | 0.31±0.01 | 0.33±0.01 | 0.08±0.02 | – | 0.91±0.02 | 91.68 | -168.09 | -179.37 |
| | A + D | 0.01±0.00 | 0.04±0.00 | 0.27±0.01 | 0.33±0.01 | 0.04±0.02 | 0.13±0.02 | 0.83±0.02 | 119.57 | -216.23 | -233.14 |
| LSA | A | 23.75±5.52 | – | 174.59±6.00 | 198.34±6.80 | 0.12±0.02 | – | 0.88±0.02 | -6503.94 | 13023.15 | 13011.88 |
| | A + D | 13.88±5.39 | 27.23±5.76 | 153.85±6.00 | 194.97±6.81 | 0.07±0.02 | 0.14±0.02 | 0.78±0.03 | -6467.27 | 12957.46 | 12940.55 |

**Note:** A, additive genetic variance; D, dominance genetic variance; E, residual variance; P, phenotypic variance; h<sup>2</sup>, additive heritability; d<sup>2</sup>, dominance ratio; e<sup>2</sup>, residual ratio; LogLik, log-likelihood; BIC, Bayesian information criterion; AIC, Akaike information criterion.

**Table 5:** Variance components, heritability (± standard error), and model fit statistics for larval body weight (LBW), length (LL), width (LW), and surface area (LSA) across diets (SYK: soy-okara, BSG: brewers’ spent grain, FVW: fruit and vegetable waste) using additive (A) and additive + dominance (A+D) models.

| Trait | Diet | Model | Variance parameters |  |  |  | Ratios |  |  | Model fit statistics |  |  |
| --- | --- | --- | --- | --- | --- | --- | --- | --- | --- | --- | --- | --- |
|  |  |  | A (Additive) | D (Dominance) | E (Residual) | P (Phenotypic) | h<sup>2</sup> (Additive) | d<sup>2</sup> (Dominance) | e<sup>2</sup> (Error) | LogLik | BIC | AIC |
| LBW | SYK | A | 399.38±70.23 | – | 685.82±40.94 | 1085.20±64.69 | 0.37±0.05 | – | 0.63±0.05 | -3757.95 | 7529.63 | 7519.90 |
| | | A + D | 276.95±63.87 | 320.86±55.09 | 420.40±35.38 | 1018.21±61.40 | 0.27±0.05 | 0.32±0.05 | 0.41±0.04 | -3691.16 | 7402.91 | 7388.32 |
| | BSG | A | 184.04±51.10 | – | 509.51±41.02 | 693.55±47.77 | 0.27±0.06 | – | 0.73±0.06 | -2058.69 | 4129.99 | 4121.39 |
| | | A + D | 80.02±44.97 | 274.33±61.87 | 316.59±40.80 | 670.94±46.79 | 0.12±0.06 | 0.41±0.08 | 0.47±0.07 | -2035.80 | 4090.52 | 4077.61 |
| | FVW | A | 99.09±31.37 | – | 364.05±28.17 | 463.15±30.73 | 0.21±0.06 | – | 0.78±0.06 | -1982.97 | 3978.58 | 3969.94 |
| | | A + D | 65.87±32.73 | 63.48±37.45 | 329.39±32.70 | 458.74±30.13 | 0.14±0.07 | 0.14±0.08 | 0.72±0.07 | -1981.11 | 3981.19 | 3968.23 |
| LL | SYK | A | 0.55±0.15 | – | 2.81±0.15 | 3.36±0.16 | 0.16±0.04 | – | 0.84±0.04 | -1059.53 | 2132.79 | 2123.06 |
| | | A + D | 0.30±0.15 | 0.67±0.17 | 2.32±0.16 | 3.29±0.17 | 0.09±0.04 | 0.20±0.05 | 0.71±0.05 | -1031.17 | 2082.91 | 2068.33 |
| | BSG | A | 0.48±0.17 | – | 2.53±0.19 | 3.03±0.19 | 0.16±0.05 | – | 0.84±0.06 | -586.93 | 1186.47 | 1177.87 |
| | | A + D | 0.38±0.19 | 0.29±0.23 | 2.33±0.22 | 3.01±0.19 | 0.13±0.06 | 0.10±0.07 | 0.77±0.07 | -577.15 | 1173.20 | 1160.30 |
| | FVW | A | 0.40±0.14 | – | 1.91±0.14 | 2.32±0.15 | 0.17±0.06 | – | 0.82±0.06 | -520.94 | 1054.53 | 1045.89 |
| | | A + D | 0.35±0.16 | 0.15±0.17 | 1.82±0.17 | 2.32±0.15 | 0.15±0.06 | 0.07±0.07 | 0.78±0.07 | -520.49 | 1059.93 | 1046.98 |
| LW | SYK | A | 0.05±0.01 | – | 0.28±0.02 | 0.33±0.02 | 0.16±0.04 | – | 0.84±0.04 | 45.03 | -76.33 | -86.05 |
| | | A + D | 0.03±0.01 | 0.07±0.02 | 0.23±0.02 | 0.32±0.02 | 0.08±0.04 | 0.22±0.05 | 0.70±0.05 | 75.01 | -129.44 | -144.03 |
| | BSG | A | 0.04±0.01 | – | 0.24±0.02 | 0.28±0.02 | 0.13±0.05 | – | 0.87±0.05 | 67.18 | -121.75 | -130.36 |
| | | A + D | 0.03±0.02 | 0.03±0.02 | 0.22±0.02 | 0.28±0.02 | 0.09±0.06 | 0.11±0.07 | 0.80±0.07 | 69.07 | -119.25 | -132.15 |
| | FVW | A | 0.03±0.01 | – | 0.25±0.02 | 0.29±0.02 | 0.11±0.05 | – | 0.89±0.05 | 60.92 | -109.21 | -117.84 |
| | | A + D | 0.03±0.02 | 0.01±0.02 | 0.25±0.02 | 0.28±0.02 | 0.09±0.06 | 0.04±0.06 | 0.87±0.06 | 61.18 | -103.41 | -116.36 |
| LSA | SYK | A | 58.99±13.63 | – | 184.43±10.53 | 243.42±13.18 | 0.24±0.04 | – | 0.76±0.05 | -3085.32 | 6184.36 | 6174.64 |
| | | A + D | 37.11±13.18 | 50.35±12.46 | 148.59±10.66 | 236.06±12.82 | 0.16±0.05 | 0.21±0.05 | 0.63±0.05 | -3062.18 | 6144.95 | 6130.36 |
| | BSG | A | 23.94±8.75 | – | 127.56±9.49 | 151.50±9.67 | 0.16±0.05 | – | 0.84±0.05 | -1656.66 | 3325.93 | 3317.33 |
| | | A + D | 11.41±9.01 | 35.87±12.58 | 102.52±10.49 | 149.80±9.72 | 0.08±0.06 | 0.24±0.08 | 0.68±0.07 | -1638.14 | 3295.18 | 3282.28 |
| | FVW | A | 19.69±6.47 | – | 73.38±5.74 | 93.08±6.20 | 0.21±0.06 | – | 0.79±0.06 | -1538.81 | 3090.26 | 3081.62 |
| | | A + D | 15.31±6.84 | 9.71±6.78 | 67.55±6.57 | 92.58±6.15 | 0.17±0.07 | 0.10±0.07 | 0.73±0.07 | -1537.32 | 3093.59 | 3080.64 |

**Note:** A, additive genetic variance; D, dominance genetic variance; E, residual variance; P, phenotypic variance; h<sup>2</sup>, additive heritability; d<sup>2</sup>, dominance ratio; e<sup>2</sup>, residual ratio; LogLik, log-likelihood; BIC, Bayesian information criterion; AIC, Akaike information criterion.

Diet-specific heritability estimates for LBW from model 2 ranged from 0.12 to 0.27, compared with an overall estimate of 0.14 (model 2). For LSA, diet-specific heritability ranged from 0.08 to 0.17, with an overall estimate of 0.07 (model 2). Heritability estimates for LL and LW were low, at 0.05 under model 2. The dominance ratio was significant across all traits and diets, indicating a consistent contribution of dominance effects. Model fit statistics indicated that model 2 performed better than model 1 in estimating the variance components.

Genetic and phenotypic correlations from model 2 for the whole dataset are given in Table 6. The genetic correlations among the growth traits were generally high and positive, with LBW and LSA showing the highest (0.99), whereas LW and LL showed relatively weak correlation (0.47). The phenotypic correlations were also positive, ranging from 0.55 to 0.83.

**Table 6:** Genetic correlations (below diagonal) and phenotypic correlations (above diagonal) (± standard error) for larval body weight (LBW), length (LL), width (LW) and surface area (LSA).

|  | LBW | LL | LW | LSA |
| --- | --- | --- | --- | --- |
| LBW | | 0.71±0.01 | 0.58±0.02 | 0.78±0.01 |
| LL | 0.84±0.08 | | 0.55±0.02 | 0.78±0.01 |
| LW | 0.82±0.10 | 0.47±0.26 | | 0.83±0.01 |
| LSA | 0.99±0.03 | 0.86±0.08 | 0.79±0.10 | |

### 3.4 Genotype by diet (G × D) interactions

G×D interactions for the four BSFL growth traits, estimated from the multi-trait additive model across the three diets, are presented in Table 7. The genetic correlation between diets ranged from -0.06 to 0.46, with all estimates well below 1, indicating varying degrees of G × D interactions. Genetic correlations between SYK and BSG were generally moderate, indicating some potential for genetic re-ranking between these diets, whereas lower correlations for some comparisons involving the FVW suggested higher potential for genetic re-ranking. For LBW, the genetic correlation was moderate between SYK and BSG (*r_g_* = 0.44 ± 0.14), but low between SYK and FVW and between BSG and FVW. A similar pattern was observed for LL, LW and LSA. Most estimates had relatively large standard errors, indicating considerable uncertainty in the magnitude of genetic correlations; however, low point estimates suggest G × D interaction and genetic re-ranking across diets. Genetic correlations were statistically significant for LBW between SYK and BSG and LL between BSG and FVW.

**Table 7:** Estimates of genotype by diet interaction measured as genetic correlation (*r_g_* ± standard error) between the three diets (SYK: soy-okara, BSG: brewers’ spent grain, FVW: fruit and vegetable waste) for larval body weight (LBW), length (LL), width (LW) and surface area (LSA).

| Trait | $r_g$ between SYK and BSG | $r_g$ between SYK and FVW | $r_g$ between BSG and FVW |
| --- | --- | --- | --- |
| LBW | 0.44±0.14* | 0.18±0.16 | 0.34±0.18 |
| LL | 0.32±0.20 | -0.06±0.21 | 0.46±0.20* |
| LW | 0.24±0.22 | 0.22±0.24 | 0.41±0.27 |
| LSA | 0.27±0.19 | 0.04±0.19 | 0.36±0.21 |

\* Significant at $p < 0.05$ (Likelihood ratio test)

## 4. Discussion

The farming of BSFL is gaining momentum worldwide, necessitating the implementation of genetic improvement programs to improve productivity, efficiency and sustainability (Gowda *et al*., 2025; Hansen *et al*., 2024). However, robust estimates of genetic parameters and G × D interactions remain limited, particularly across genetically distinct populations and diverse production environments under which it is farmed. These parameters are critical for designing effective breeding strategies, predicting the potential for genetic improvement, and informing strain selection and management in practical BSFL production systems. Optimising strains under specific breeding or test environments may not necessarily translate to different rearing conditions; therefore, understanding the extent of G × D is crucial. In this study, we present the first estimates of genetic parameters and G × D interactions for the Australian BSFL population, representing the Australasian genetic lineage, which exhibits clear genetic differentiation (i.e., diversity and divergence) from other global populations (Generalovic *et al*., 2025b; Kaya *et al*., 2021). By genotyping a genome-wide panel of 6K SNPs across 2,083 individuals, we provide the first comprehensive integration of genomic and phenotypic data to quantify heritable variation and diet-specific genetic responses in this lineage. These findings establish a baseline for breeding in this lineage and provide practical insights for optimising larval performance across production environments.

### 4.1 Non-genetic factors

The present study encompassed three diets (SYK, BSG, and FVW), which strongly influenced growth traits in BSFL. Most of the variation in larval growth traits was attributable to diet, consistent with the nutritional composition of the substrates, which largely determines larval performance (Danieli *et al*., 2019; Meneguz *et al*., 2018; Scala *et al*., 2020). Although SYK and BSG contained similar levels of crude protein (20-21%) (supplementary Table 1), larval growth performance on these diets differed, suggesting that crude protein content alone does not explain the observed differences. These differences may be attributable to variation in protein quality, particularly the balance of essential amino acids and overall digestibility (Bava *et al*., 2019; Chia *et al*., 2020; Danieli *et al*., 2019). Additionally, unmeasured differences in fibre content and the physical properties of substrates, such as texture and palatability, may also have influenced variation in larval growth (Chia *et al*., 2020). As both diets are secondary by-products, the processing methods themselves are relevant: BSG is typically milled, whereas SYK is boiled during milk extraction. Such differences in processing may alter nutrient availability and absorption in BSFL. In addition, microbial load in the feed substrates may affect gut health and thereby influence growth performance (Hopkins *et al*., 2021). In the case of FVW, its low nutritional profile, combined with high fibre content, resulted in consistently poor larval growth (Meneguz *et al*., 2018). Optimising BSFL feed by mixing these substrates in appropriate proportions may therefore enhance growth performance and improve the utilisation of diverse organic waste sources (Oonincx *et al*., 2015).

We evaluated larval colour as a phenotypic proxy for the transition between life stages; it influenced all traits studied. Larvae progressively darken after crossing the fifth instar, making colour a reliable indicator of developmental/harvest stage (Bruno *et al*., 2025). As larvae darkened, body weight began to decrease, suggesting that pigmentation reflects both developmental progression and underlying physiological differences (Liu *et al*., 2017). Thus, larval colour can help identify the optimal harvest timing, with the fifth instar yielding a superior final product in terms of nutritional quality. Including colour as a covariate improved the model by capturing the growth dynamics and stage-related variation. With a better understanding of these mechanisms, larval colour could also be considered a trait for selection, indirectly assisting in identifying developmental stage and improving breeding and production strategies.

Given the short BSFL grow-out period (∼7 days), the sampling day significantly affected growth traits. On sampling day 2, the least squares mean for LBW was highest (138.45 ± 1.17 mg). As larvae progressed from the fourth to the fifth instar, growth rates changed markedly. After the fifth instar, larvae ceased feeding, began to lose weight, and exhibited darker pigmentation. These changes indicate underlying physiological and metabolic differences, and it is evident that nutrient assimilation also varies across developmental stages.

Each tray served as a dynamic microenvironment, functioning as an independent unit whose interactions with the diet influenced larval growth. Although trays within diets were maintained under uniform housing conditions, including temperature, relative humidity, and consistent substrate quantity and larval density, microenvironmental variation persisted. This highlights the need to carefully consider and optimise tray setup. Larval activity within trays further affected performance, as larvae compete for and assimilate nutrients differently. Mortality rates between trays also contributed to variation in final yield, emphasising the importance of minimising tray-to-tray differences to ensure reliable growth outcomes.

### 4.2 Genetic parameters

In this study, we estimated genetic parameters for growth traits in the Australasian BSFL population, providing a baseline for implementing a selective breeding program. Pedigree information and parental genotypes were not available, as it is difficult to tag and track family structure in BSFL commercial production systems (Slagboom *et al*., 2024). Eggs were hatched in a single nursery environment to reduce common environmental effects, and SNPs were used to establish accurate relationships among individuals. This approach enabled estimation of genetic parameters with a sufficient sample size (n = 2083). Because the experimental design included three diets, heritability was estimated both diet-specific and overall. The major source of variation in growth traits was attributable to diet.

To capture the full genetic architecture of traits, we initially fitted a standard additive genetic model. However, given the crossbred nature of the population, we also tested an additive– dominance model, which provided a better fit (Tables 4 and 5). Overall heritability estimates were low, ranging from 0.04 ± 0.02 to 0.14 ± 0.03 across all diets. LBW showed low heritability (0.14 ± 0.03) due to a significant dominance effect (0.19 ± 0.02), which reduced the heritability from 0.20 in the additive-only model to 0.14 in the additive-dominance model. The inclusion of dominance effects was significant, indicating that non-additive genetic variation plays an important role in BSFL growth traits, which is supported by the earlier study (Meyermans *et al*., 2025). However, this should be interpreted cautiously because dominance variance can potentially absorb shared environmental effects in family-based designs.

Diet-specific heritability estimates ranged from 0.08 ± 0.04 to 0.37 ± 0.05. These values were consistently higher than the overall estimates, suggesting that expression of additive genetic variance and its relative contribution to phenotypic variation differed among diets. Based on these estimates, selective breeding to improve LBW is feasible; they are similar to the previous study (Zaalberg *et al*., 2025). Genetic correlations among growth traits were positive and high, ranging from 0.47 to 0.99 (model 2), and were consistent with phenotypic correlations. As reported in previous studies, the strong genetic correlation between LBW and LSA suggests that LSA could be used as an indirect selection criterion, particularly when implementing computer vision systems, which could be efficient in insect production (Generalovic *et al*., 2025a). The presence of significant dominance effects in the current population, likely due to crossbred stocks and the polyandrous mating system of BSFL (Hoffmann *et al*., 2021). This further suggests that crossbreeding strategies could be pursued to exploit heterosis (hybrid vigour) and thereby enhance productivity in BSFL farming.

### 4.3 Genotype by diet (G × D) interactions

The present population exhibited moderate G × D interactions (Table 7). Variance estimates from the multivariate additive–dominance model were constrained at the boundary and failed to converge, possibly due to the relatively small sample sizes in BSG (n=555) and FVW (n=563) compared with SYK (n=965). Consequently, an additive multivariate model was fitted. The genetic correlation between LBW in SYK and BSG was 0.44, whereas genetic correlations across the other diet combinations ranged from near zero to moderate values, indicating greater potential for G × D interactions and genetic re-ranking, particularly for comparisons involving FVW. The non-significant genetic correlation estimates should not be interpreted as evidence for the absence of G × D interactions, as their relatively large standard errors indicate considerable uncertainty in these estimates. This discrepancy in sample sizes may have contributed to the difficulty in obtaining accurate variance estimates. Furthermore, in the absence of confirmed parent-offspring family structure information, it is challenging to determine whether individuals within each family were evenly distributed across diets. Although *COLONY* (Jones and Wang, 2010) inferred 64 family clusters, 27 had sufficient representation across diets (≥5 individuals in each diet), including 18 clusters with ≥10 individuals in each diet (Supplementary Table 4). Examination of the top 5% of individuals ranked by GEBVs and visualisation of their reaction norms (Figure 3) further indicated genetic re-ranking across diets. The extensive genetic re-ranking observed for LL, particularly between SYK and FVW, was consistent with the very low genetic correlation between these diets (−0.06 ± 0.21), whereas the higher correlations for the other diet pairs indicated greater consistency in genetic ranking. Previous studies have indicated the presence of G × D interactions in BSFL, with the magnitude varying between populations (Generalovic *et al*., 2025a; Sandrock *et al*., 2022). In this study, we detected G × D interactions in the Australian BSFL population, which has important implications for how this information can be incorporated into breeding programs. More balanced sample sizes, structured family design with families represented across diets and greater replication across trays and genetic backgrounds would likely improve the precision of genetic correlation estimates and provide stronger evidence for the magnitude of G × D interactions.

**Figure 3:**
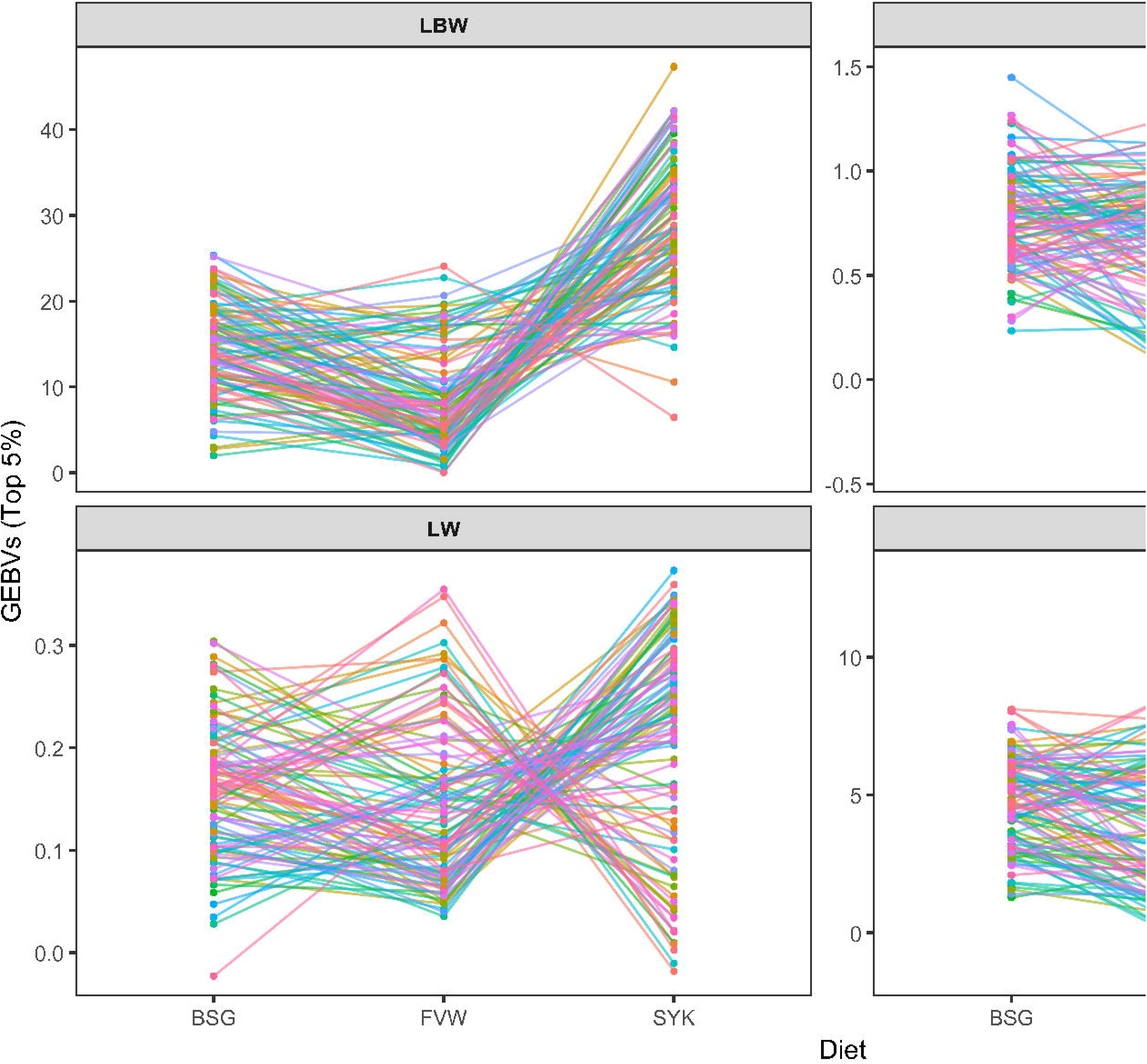
Reaction norms of top 5% individuals illustrating re-ranking of genomic estimated breeding values (GEBVs) across diets (SYK: soy-okara, BSG: brewers’ spent grain, FVW: fruit and vegetable waste) for larval body weight (LBW), length (LL), width (LW) and surface area (LSA). Each coloured line represents an individual larva, demonstrating moderate genotype-by-diet (G×D) interactions.

### 4.4 Implications for breeding programmes

The estimated heritabilities for LBW and LSA in BSFL indicate a moderate genetic basis for the traits, which can be exploited in selective breeding programs to improve growth performance. Given the short life cycle of BSF (∼38-45 days) and the moderate heritability of these traits, sustained genetic gain through selection is feasible. Although recent studies have demonstrated the feasibility of producing full-sib and half-sib families and tracking pedigree information (Jensen *et al*., 2025; Zaalberg *et al*., 2025), implementing family-based selection or controlled mating schemes remains challenging in commercial production systems, given the large census population sizes. In this context, line breeding represents a more practical alternative, as it requires fewer resources and can be readily implemented under mass-rearing conditions. Importantly, the current results indicate that G × D interactions are non-negligible and must be considered when designing selective breeding strategies. The low to moderate genetic correlations among diets suggest that selection under one diet may not necessarily translate into equivalent genetic gain under another. Therefore, breeding programmes should consider whether to target strains adapted to a specific standardised diet or strains with more stable performance across production environments, with the breeding environment aligned with the intended commercial rearing conditions, as is commonly considered in plant breeding. Given the magnitude of the G × D interactions observed here, ignoring these interactions could reduce the transferability of genetic gain from the breeding environment to the commercial production environment, as highlighted by previous simulation work in BSF (Slagboom *et al*., 2024). In addition, heterosis can be exploited through controlled crosses among genetically distinct lines, thereby enhancing performance, mitigating inbreeding, and helping maintain long-term genetic diversity within large production systems (Windig and Kaal, 2008).

Looking ahead, the continued development of genomic resources, particularly rapid and cost-effective genotyping platforms and fast-track sequencing technologies such as Nanopore, is likely to make genomics-assisted selection strategies feasible in BSFL, thereby accelerating genetic improvement. Although non-invasive DNA extraction and individual tracking across metamorphosis remain challenging under the current production systems, especially given the short life cycle, ongoing technological advances suggest that the practical implementation of genomics-assisted selection could become achievable in the near future. Individual-based GEBVs remain impractical in mass-rearing systems; instead, genomic prediction is more likely to be applied to rank breeding lines and select elite lines. Furthermore, genomic information could be strategically used to monitor inbreeding levels, introduce external germplasm into the breeding nucleus (provided sufficient connectivity to the training population, for example, through common ancestry), and ensure reliable prediction, thereby maintaining both genetic merit and diversity.

## 5. Conclusions

We found considerable effects of diet, sampling day, and larval colour on BSFL growth traits, highlighting the importance of refining management practices to optimise production. The diet-specific heritability estimates indicate that traits such as LBW or LSA can be considered as selection criterion to improve growth performance. However, this study was limited to growth-related traits; other product quality traits such as protein and fat content, developmental time, and reproductive efficiency should be investigated in future research. Given the evidence of G × D interactions in the present population, breeding programs need to be tailored to account for these effects. The substantial dominance effects observed suggest that the initial generations of the assembled base population still carry non-additive genetic variation, which could be exploited in breeding strategies.

## Acknowledgements

We acknowledge the Australian Research Council for funding through a Linkage Project (LP210301250), FlyFarm Australia Ltd for providing facilities for this study, and Zina Patel for assistance in managing the BSF colonies. We also thank Jessica Maddams and Xinyue Cui for their assistance with phenotyping.

## Author contributions

Kishor B. Gowda (Conceptualization, Data curation, Formal analysis, Investigation, Methodology, Writing – original draft), Shafira Septriani (Investigation, Writing – review & editing), David B. Jones (Resources, Writing – review & editing), Dean R. Jerry (Conceptualization, Funding acquisition, Writing – review & editing), Constant Tedder (Funding acquisition, Resources, Writing – review & editing) and Kyall R. Zenger (Conceptualization, Funding acquisition, Investigation, Project administration, Resources, Supervision, Writing – review & editing)

## Data availability

The phenotypic and genotypic data have been deposited in the <u>JCU Research Data</u> repository and will be made publicly accessible upon acceptance of the manuscript. The scripts used for the analyses are available at https://github.com/KB-Gowda/BSF_Quantitative_Genomics.git.

