## Supplementary for "Genetic parameters and genotype-by-diet interactions for growth traits in Australian black soldier fly larvae: Implications for selective breeding"

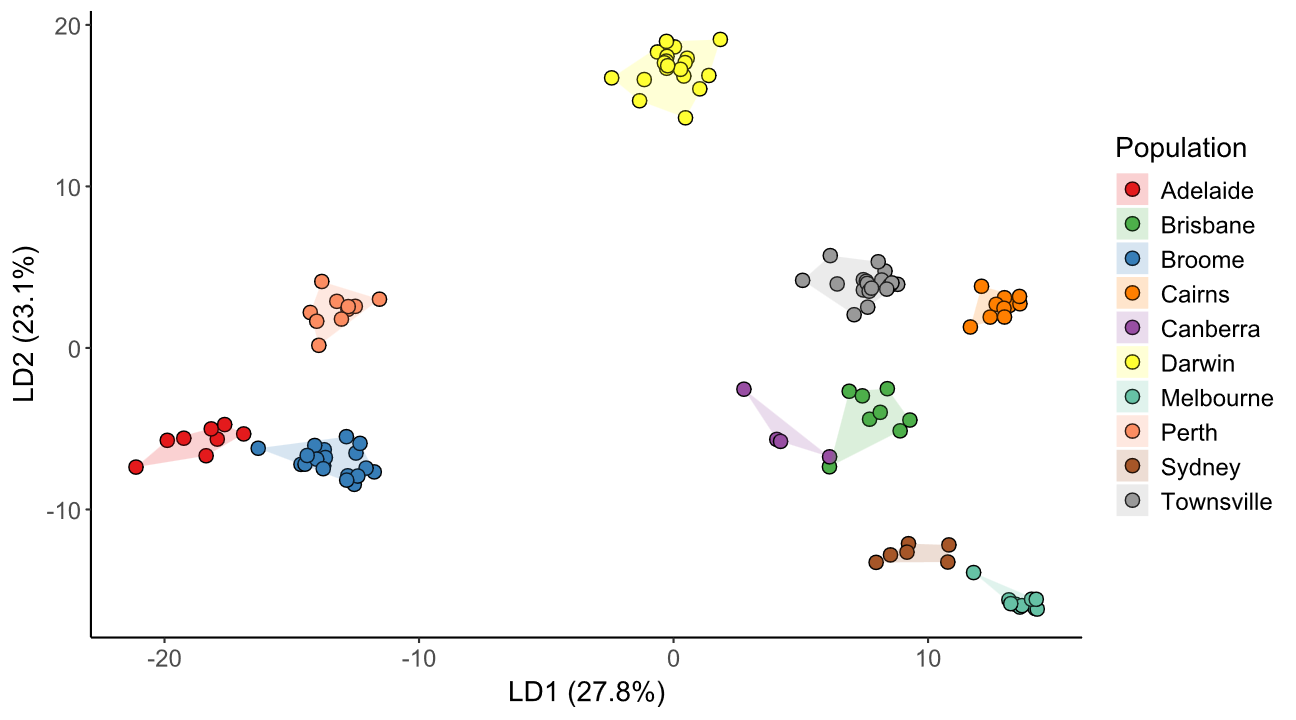

**S_Fig 1**: Discriminant analysis of principal components (DAPC) showing genetic differentiation among BSF populations collected across Australia

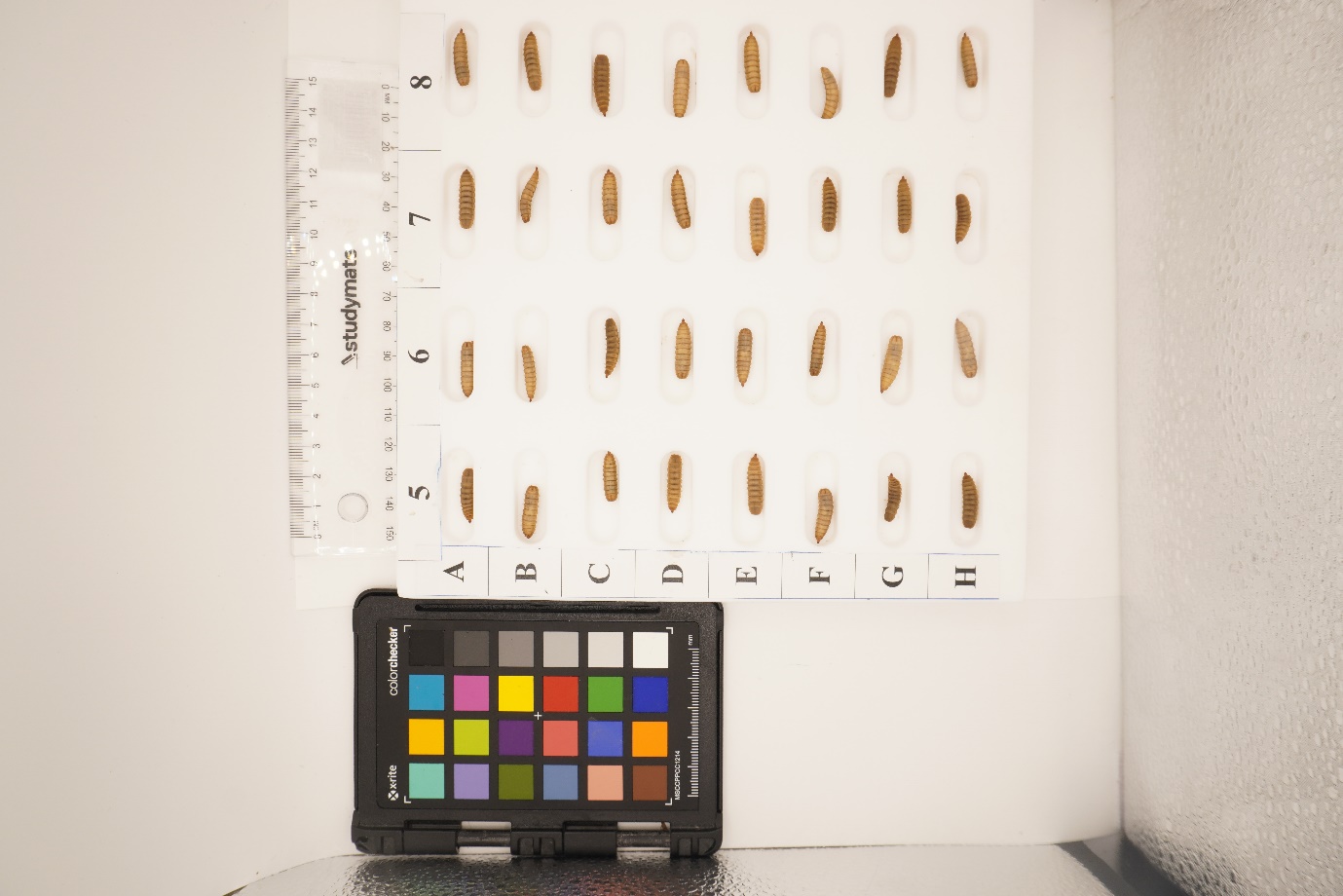

**S_Fig. 2**: Capture of BSFL images by placing individual larvae in the grooves carved into a teflon board

**S_Table 1**: Proximate composition of BSFL rearing substrates

| **Component** | **FVW** | **SYK** | **BSG** |
| --- | --- | --- | --- |
| Lipid (%) | 1.23 | 4.24 | 9.37 |
| Protein (%) | 7.30 | 21.30 | 20.25 |
| Ash (%) | 4.03 | 3.75 | 4.28 |

**S_Table 2**: Number of BSFL samples across trays, diets and sampling days

| **Tray** | **Day 13** | | | **Day 14** | | | **Day 15** | | |
| --- | --- | --- | --- | --- | --- | --- | --- | --- | --- |
|  | **BSG** | **SYK** | **FVW** | **BSG** | **SYK** | **FVW** | **BSG** | **SYK** | **FVW** |
| **T1** | 94 | 128 | 94 | 48 | 110 | 56 | 61 | 95 | 63 |
| **T2** | 92 | 125 | 48 | 46 | 109 | 70 | 62 | 100 | 62 |
| **T3** | 48 | 128 | 46 | 46 | 119 | 62 | 63 | 59 | 63 |
| **Total** | 234 | 381 | 188 | 140 | 338 | 188 | 186 | 254 | 188 |
| **Overall** | **803** | | | **666** | | | **628** | | |

**S_Table 3**: Descriptive statistics of BSFL growth traits (LBW, LL, LW and LSA) across diets and overall (including the number of observations, minimum and maximum values, mean with standard error, standard deviation, and coefficient of variation)

| **Diet** | **Traits** | **N** | **Min** | **Max** | **Mean (SE)** | **SD** | **CV (%)** |
| --- | --- | --- | --- | --- | --- | --- | --- |
| **SYK** | **LBW (mg)** | 973 | 55 | 266 | 177.11 (1.15) | 35.97 | 20.31 |
|  | **LL (mm)** | 973 | 12.33 | 23.89 | 19.02 (0.07) | 2.05 | 10.77 |
|  | **LW (mm)** | 973 | 2.61 | 6.28 | 4.87 (0.01) | 0.62 | 12.77 |
|  | **LSA (mm^2^)** | 973 | 35.01 | 130.24 | 79.60 (0.59) | 18.49 | 23.24 |
| **BSG** | **LBW (mg)** | 560 | 62 | 218 | 141.85 (1.12) | 26.61 | 18.76 |
|  | **LL (mm)** | 560 | 11.38 | 22.73 | 17.80 (0.07) | 1.75 | 9.84 |
|  | **LW (mm)** | 560 | 2.95 | 5.79 | 4.57 (0.02) | 0.53 | 11.61 |
|  | **LSA (mm^2^)** | 560 | 32.95 | 101.75 | 69.49 (0.52) | 12.49 | 17.97 |
| **FVW** | **LBW (mg)** | 564 | 10 | 167 | 81.82 (1.06) | 25.34 | 30.96 |
|  | **LL (mm)** | 564 | 9.09 | 18.68 | 14.17 (0.07) | 1.75 | 12.33 |
|  | **LW (mm)** | 564 | 1.31 | 5.41 | 3.97 (0.02) | 0.65 | 16.40 |
|  | **LSA (mm^2^)** | 564 | 11.99 | 85.85 | 47.58 (0.54) | 12.86 | 27.04 |
| **Overall** | **LBW (mg)** | 2097 | 10 | 266 | 142.07 (1.09) | 50.07 | 35.24 |
|  | **LL (mm)** | 2097 | 9.09 | 23.89 | 17.39 (0.06) | 2.76 | 15.89 |
|  | **LW (mm)** | 2097 | 1.31 | 6.28 | 4.55 (0.01) | 0.71 | 15.6 |
|  | **LSA (mm^2^)** | 2097 | 11.99 | 130.24 | 68.29 (0.44) | 20.49 | 30 |

**S_Table 4**: Distribution of individuals across the three experimental diets (BSG, SYK, and FVW) within each of the 69 identified colony clusters.

| **Cluster** | **BSG** | **SYK** | **FVW** | **Total** |
| --- | --- | --- | --- | --- |
| 1 | 25 | 25 | 26 | 76 |
| 2 | 22 | 51 | 35 | 108 |
| 3 | 37 | 53 | 35 | 125 |
| 4 | 5 | 24 | 10 | 39 |
| 5 | 13 | 30 | 17 | 60 |
| 6 | 3 | 7 | 6 | 16 |
| 7 | 40 | 64 | 45 | 149 |
| 8 | 13 | 30 | 19 | 62 |
| 9 | 4 | 10 | 4 | 18 |
| 10 | 20 | 27 | 18 | 65 |
| 11 | 22 | 33 | 20 | 75 |
| 12 | 7 | 22 | 18 | 47 |
| 13 | 41 | 53 | 31 | 125 |
| 14 | 27 | 44 | 34 | 105 |
| 15 | 38 | 49 | 22 | 109 |
| 16 | 26 | 46 | 29 | 101 |
| 17 | 15 | 33 | 21 | 69 |
| 18 | 6 | 13 | 6 | 25 |
| 19 | 20 | 28 | 20 | 68 |
| 20 | 22 | 22 | 8 | 52 |
| 21 | 15 | 35 | 12 | 62 |
| 22 | 15 | 16 | 16 | 47 |
| 23 | 1 |  |  | 1 |
| 24 | 10 | 21 | 5 | 36 |
| 25 | 1 |  |  | 1 |
| 26 | 10 | 15 | 6 | 31 |
| 27 | 10 | 21 | 10 | 41 |
| 28 | 17 | 22 | 18 | 57 |
| 29 | 4 | 11 | 3 | 18 |
| 30 |  | 7 | 4 | 11 |
| 31 |  | 3 |  | 3 |
| 32 | 8 | 9 | 6 | 23 |
| 33 | 4 | 11 | 5 | 20 |
| 34 | 6 | 7 |  | 13 |
| 35 | 7 | 7 | 2 | 16 |
| 36 | 7 | 19 | 4 | 30 |
| 37 | 2 | 9 | 2 | 13 |
| 38 | 4 | 11 | 1 | 16 |
| 39 | 4 | 12 |  | 16 |
| 40 | 1 | 9 | 1 | 11 |
| 41 |  | 1 |  | 1 |
| 42 | 5 | 11 | 7 | 23 |
| 43 | 2 | 7 |  | 9 |
| 44 |  | 1 |  | 1 |
| 45 | 7 | 8 | 7 | 22 |
| 46 |  |  | 1 | 1 |
| 47 | 3 | 3 | 12 | 18 |
| 48 | 2 | 2 | 1 | 5 |
| 49 | 2 | 14 | 5 | 21 |
| 50 |  | 3 | 2 | 5 |
| 51 |  |  | 3 | 3 |
| 56 | 1 |  |  | 1 |
| 57 | 1 |  |  | 1 |
| 58 | 3 | 1 | 2 | 6 |
| 59 |  | 1 |  | 1 |
| 60 |  | 2 |  | 2 |
| 61 | 1 | 3 | 1 | 5 |
| 62 |  | 1 |  | 1 |
| 63 |  | 1 |  | 1 |
| 64 |  | 1 |  | 1 |
| 65 |  | 1 |  | 1 |
| 66 |  |  | 1 | 1 |
| 67 |  |  | 1 | 1 |
| 68 |  |  | 1 | 1 |
| Grand Total | 559 | 970 | 563 | 2092 |
